# Cpf1(Cas12a)-based genome editing in the filamentous cyanobacterium *Nostoc punctiforme*

**DOI:** 10.64898/2026.07.31.742098

**Authors:** Jenna R. Ryder, Soohan Woo, Ailea A. Blahm, Christine N. Cummings, Douglas D. Risser

**Affiliations:** Department of Biology, University of Colorado Colorado Springs, Colorado Springs, CO 80918 USA; Department of Biology, University of the Pacific, Stockton, CA 95211 USA

**Keywords:** Cyanobacteria, CRISPR, *N. punctiforme*, *Cpf1*

## Abstract

The filamentous cyanobacterium *Nostoc punctiforme* is a key model organism used to study several aspects of cyanobacterial biology, including development, nitrogen-fixing symbioses with plants, and secondary metabolites, among others. While *N. punctiforme* is amenable to genetic manipulation, traditional approaches for the generation of mutant strains using homologous recombination are slow, requiring prolonged outgrowth under antibiotic selection to ensure isogenic mutant populations. CRISPR-based genome editing using Cpf1 (Cas12a) was recently shown to be an effective means of rapid generation of isogenic mutants in several cyanobacteria. In this study, Cpf1-based genome editing tools were developed for *N. punctiforme*. A total of 19 unmarked, in-frame deletion mutants were successfully constructed using Cpf1-targeted cleavage along with homology directed repair (HDR). The length of the homology arms (HAs) on the homologous repair template (HRT) used for HDR was found to be a critical factor for successful deletion of target genes, with some requiring up to 4 kb HAs to acquire mutant exconjugants. A strategy for allelic replacement was also developed by introducing an exogenous target site in place of the deleted genes, which could subsequently be targeted for cleavage and repaired with an HRT containing altered alleles of the genes of interest. Additionally, a single-step cloning strategy was devised, allowing for rapid assembly of editing plasmids, and improved conjugation protocols for genetic transfer from *E. coli* to *N. punctiforme* were implemented. Collectively, these tools and protocols should enhance the pace and ease of conducting genetic studies in this important model cyanobacterium.

## Introduction

Cyanobacteria are distinguished by their ability to perform oxygenic photosynthesis, and range from simple unicellular forms to complex, multicellular species that exhibit substantial developmental complexity (Rippka et al., 1979). *Nostoc punctiforme* ATCC 29133 (syn. PCC 73102) is an example of the latter that serves as a critical model organism because it is genetically tractable and exhibits a wide variety of phenotypes observed in multicellular cyanobacteria (Risser, 2023). *N. punctiforme* grows as unbranched filaments of vegetative cells that perform oxygenic photosynthesis, but it can differentiate various cell/filament types, including nitrogen-fixing heterocysts, motile hormogonia, and environmentally resistant resting cells termed akinetes (Risser, 2023). Notably, while the molecular regulation of heterocyst development has been more intensively studied in the widely adopted model filamentous cyanobacterium *Nostoc* sp. PCC 7120 (Zeng and Zhang, 2022), this cyanobacterium does not exhibit hormogonia or akinete differentiation, making *N. punctiforme* a critical system to study these developmental pathways (Risser, 2023).

In addition to its utility in understanding multicellular development, *N. punctiforme* is a critical organism for the study of plant-cyanobacterial symbiosis, the production of secondary metabolites, and the formation of biofilms and aggregate structures known as photogranules. Originally isolated from the coralloid root of a cycad (Meeks et al., 2001), *N. punctiforme* can establish nitrogen-fixing symbioses with numerous plant hosts and is currently the only filamentous cyanobacterium capable of forming these associations that is amenable to genetic manipulation (Álvarez et al., 2023). A better understanding of the molecular basis for these interactions could potentially lead to leveraging cyanobacteria as biofertilizers in engineered symbioses with crop plants (Álvarez et al., 2023). Secondary metabolites investigated in *N. punctiforme* include the sunscreen pigments scytonemin (Ferreira and Garcia-Pichel, 2016; Klicki et al., 2018; Klicki et al., 2022; Naurin et al., 2016; Parrett et al., 2025; Soule et al., 2007) and shinorine (Balskus and Walsh, 2010; Gao and Garcia-Pichel, 2011), the halogenated nostochlorosides (Glasser et al., 2024), and various other polyketide synthase (PKS) (Liaimer et al., 2011) and non-ribosomal peptide synthase (NRPS) derived compounds (Dehm et al., 2019), such as nostopeptolide (Liaimera et al., 2015), which plays a key signaling role in hormogonium development. Understanding the molecular foundations of scytonemin biosynthesis is of particular interest, as this molecule has potential cosmetic and medical applications (Behera et al., 2025). Photogranules are macroscopic, spherical aggregates formed by filamentous cyanobacteria that represent a surface-detached biofilm (Abouhend et al., 2020). Their formation appears to be driven by the acquisition of motility, and mutations that abolish hormogonium development and motility eliminate photogranulation (Parrett et al., 2025; Risser, 2025). Photogranules are of considerable interest for applications in wastewater treatment (Wang et al., 2026), and an understanding of the molecular drivers of this phenotype may be essential for artificial control in these settings.

Currently, techniques for genomic manipulation of *N. punctiforme* rely on homologous recombination, and our research group has routinely used this approach to generate unmarked, in-frame gene deletions (for example, see (Khayatan et al., 2015; Risser et al., 2014)). However, despite the high rate of success, this approach is slow due to three factors: 1) the slower growth rate of *N. punctiforme* compared to unicellular model cyanobacteria, 2) the requirement for prolonged outgrowth of single-recombinants on antibiotic selection to ensure isogenic populations prior to counterselection of double-recombinants, likely due to the polyploid nature of cyanobacteria (Griese et al., 2011), and 3) the need to sometimes screen large numbers of double-recombinant colonies (10s-100s) via PCR to identify those with the genotype of interest, rather than those that have reverted to the wild type. Therefore, new strategies that enhance the rate of genetic manipulation in *N. punctiforme* have the potential to reduce this bottleneck and thus advance the productivity of research in this organism.

One promising approach is the implementation of CRISPR-based genome editing. While the CRISPR/Cas9 system (Ran et al., 2013), commonly employed for genome editing in many organisms, has not been widely adopted in cyanobacteria due to apparent toxicity of the Cas9 nuclease (Ungerer and Pakrasi, 2016), the more recently developed Cpf1 (Cas12a) system (Zetsche et al., 2015) has been utilized successfully in several cyanobacteria (Niu et al., 2019; Ungerer and Pakrasi, 2016). Precise genome editing is typically achieved by introducing a replicative plasmid expressing the *cpf1* gene and CRISPR array, with one of the spacers in the array modified to produce a crRNA (*i.e.* gRNA) which, once expressed and processed, will complex with Cpf1 and target the gene of interest (GOI) for cleavage. In addition, this plasmid contains a homologous repair template (HRT), which in the case of a gene deletion, will consist of homology arms (HAs) with DNA corresponding to the flanking 5’ and 3’ regions of the GOI, fused together. Following cleavage by Cpf1, homology directed repair (HDR) can utilize this HRT to fix the lesion, resulting in deletion of the target gene. Importantly, because Cpf1 cuts DNA 3’ and distal to the PAM site (Zetsche et al., 2015), non-homologous end joining (NHEJ) may not erase the target site for the gRNA and thus allow repeated cleavage of the same genomic region. Therefore, this system should favor HDR over NHEJ when cells are supplied with an HRT that facilitates removal of the target gene. The editing plasmid can subsequently be cured by outgrowth without antibiotic selection (Ungerer and Pakrasi, 2016), and in some cases, the inclusion of a counter-selectable marker such as *sacB* on the plasmid (Niu et al., 2019).

In this study, we demonstrate successful implementation of Cpf1-based gene deletion and allelic replacement in *N. punctiforme*, identifying HA length as a critical limiting factor for successful editing of some genes. Additionally, we develop a streamlined, single-step cloning strategy for rapid construction of editing plasmids with CRISPR arrays containing multiple gRNAs, and an optimized, user-friendly conjugation protocol for transfer of editing plasmids from *E. coli* to *N. punctiforme*. Collectively, implementation of these tools and protocols has the potential to substantially enhance the rate of genetic studies in *N. punctiforme*.

## Results

### Homologous repair template (HRT) length is a critical factor in successful editing

To test Cpf1-based gene deletion in *N. punctiforme*, three genes, *hmpF*, *hfq*, and *pilB*, were initially targeted. These genes were chosen because deletion mutants have previously been constructed using homologous recombination, and they are easy to assess phenotypically due to their loss of motility (Cho et al., 2017; Harwood et al., 2021; Khayatan et al., 2015). The plasmid targeting *hmpF* was built from plasmid pCpf1b as previously described (Niu et al., 2019), with ∼1kb each of 5’ and 3’ flanking DNA for the HRT and incorporating a single gRNA targeting *hmpF*, into the CRISPR array. However, we found this cloning strategy to be somewhat cumbersome, as it requires the sequential construction of 2 plasmids, the first containing the gRNA, which is then used to assemble a second plasmid including the HRT. For the plasmids targeting *hfq* and *pilB*, a single-step cloning strategy was developed where the CRISPR array with the gene-specific gRNA was generated by gene synthesis (Fig. S1), and the array, 5’ and 3’ HAs (∼1kb each) were amplified via PCR and fused with linearized pCpf1b digested with AvrII and BamHI using a Gibson-based (Gibson et al., 2009) (HiFi) assembly strategy (Fig. 1, see materials and methods for further details). These plasmids were subsequently transferred from the donor *E. coli* strain UC585 (Liang et al., 1993) into *N. punctiforme* via biparental mating using an optimized conjugation protocol (see materials and methods for details).

**Figure 1.**
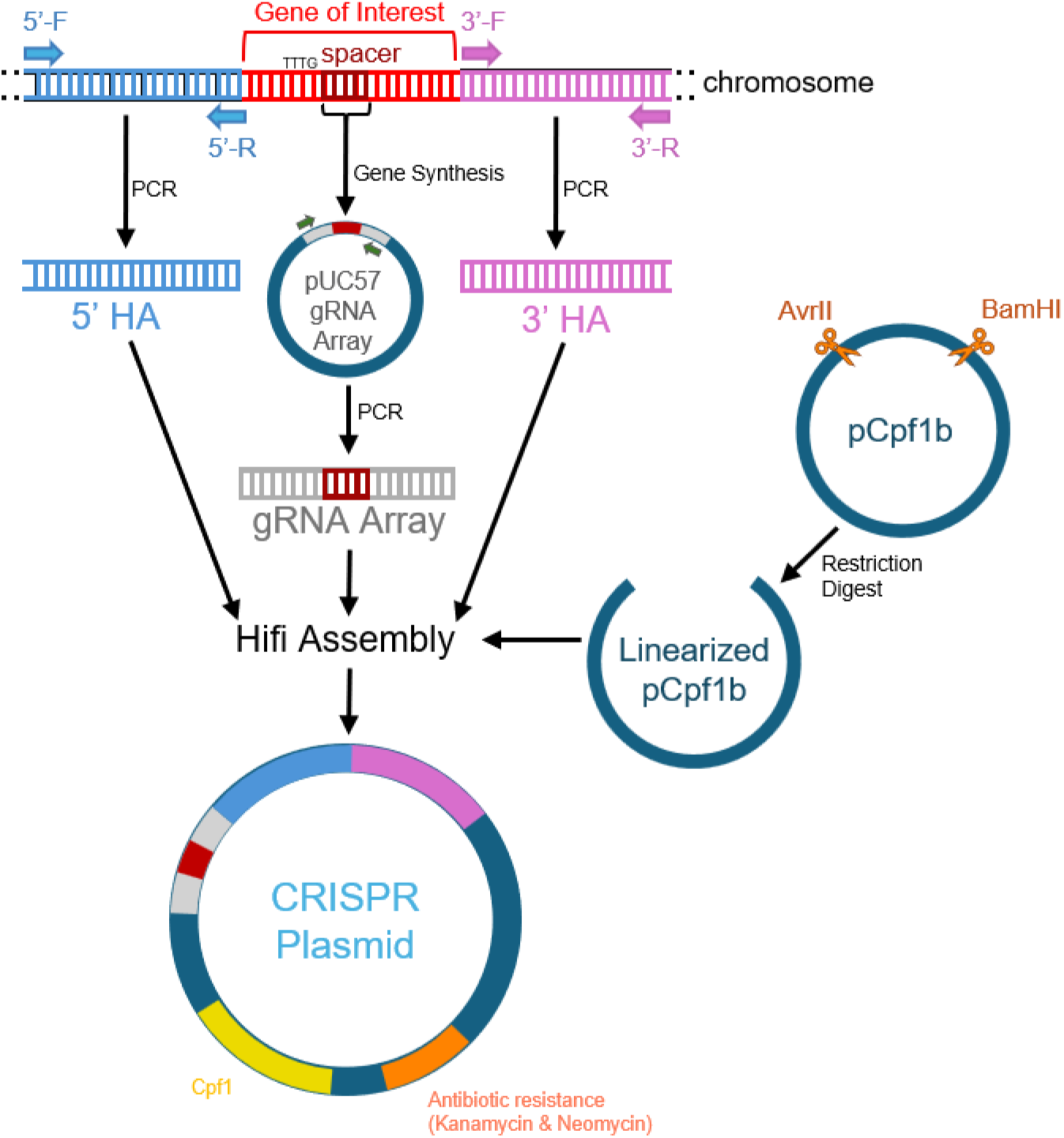
Schematic diagram of editing plasmid assembly. One or two 22 bp gRNAs immediately adjacent to TTTG PAM sites are selected for the gene of interest and a CRISPR array with the gRNA/s is synthesized and cloned into pUC57 (see Fig. S1 for example gRNA array sequences). The gRNA array and ∼1-4 kb 5’ and 3’ HAs are amplified by PCR and then HiFi assembled into plasmid pCpf1b linearized by digestion with AvrII and BamHI.

The plasmids targeting *hmpF* and *hfq* produced hundreds of exconjugants, with genotypic and phenotypic tests indicating that 100% of these colonies were fully segregated deletion mutants (Fig. 2A-C, Fig. S2). However, the plasmid targeting *pilB* failed to produce any exconjugants. There are several reasons why this may be the case: 1) there might be a toxic gene on the *pilB* HRT, 2) there is off-target cutting of the gRNA, which is lethal, and 3) there could be inefficient HDR following cleavage of *pilB*, which is lethal either because NHEJ is not sufficiently active to repair the lesion or does not remove the target site and thus allows repeated cleavage of the chromosome. To distinguish between these possibilities, the plasmid targeting *pilB* was introduced into a previously generated Δ*pilB* strain (Khayatan et al., 2015). If there is a toxic gene in the HRT or off-target cutting in the gRNA, then the plasmid should not produce exconjugants in either the wild type or the *pilB* mutant. In contrast, if there is inefficient homologous repair of *pilB*, then the plasmid should produce exconjugants in the Δ*pilB* strain because the target site for the gRNA is absent. Unlike the wild type, introduction of this plasmid into the Δ*pilB* mutant produced many exconjugants, indicating that the third possibility is the most likely (Fig. S3).

**Figure 2.**
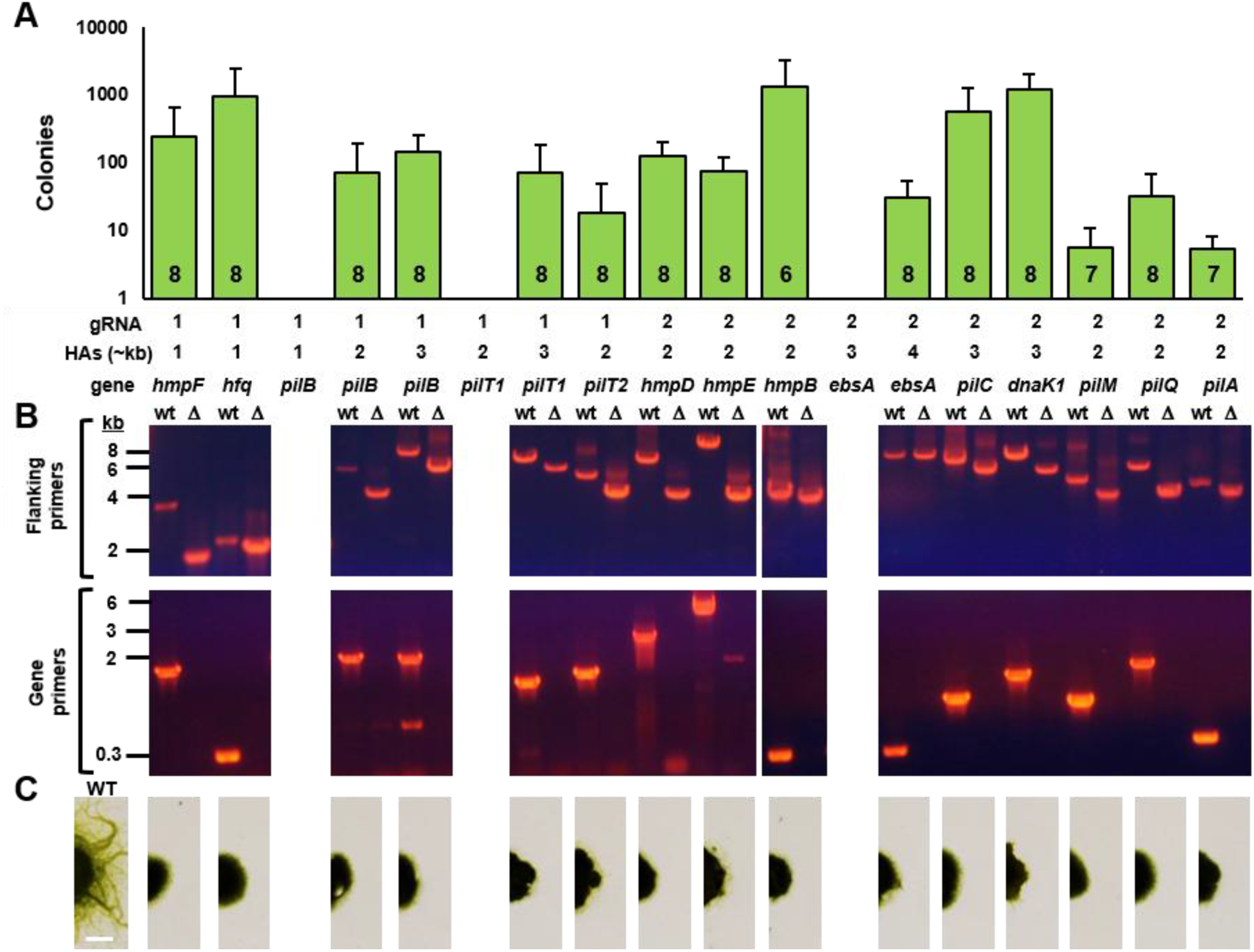
Cpf1-based deletion of motility genes in *N. punctiforme*. (**A**) Average number of exconjugants from three independent conjugations for each plasmid tested. Error bar = 1 S.D. Number of gRNAs, size of HAs, and target gene for each plasmid are indicated. Number inside each bar on graph is the number of colonies, out of 8 tested, that displayed the mutant phenotype. (**B**) PCR-based genotype analysis of mutants constructed for plasmids that yielded exconjugants. PCR reactions either utilized primers that anneal just outside of the region used for the HRT (flanking primers), and produce a smaller amplicon compared to the wild type if the GOI is deleted, or primers that anneal to the beginning and end of the target gene (gene primers) and produce no amplicon if the GOI is deleted (see figure S2 for a schematic diagram of primer binding sites). Identity of target genes corresponds to labels in panel A. (**C**) Phenotypic analysis of motility in the mutant strains by colony spreading assays. The wild type strain exhibits colony spreading while the mutant strains exhibit abolished or severely diminished colony spreading. White bar = 0.5 mm. Identity of mutant strains corresponds to labels in panel A.

The efficiency of HDR is known to be positively correlated with the length of the homology arms (Chai et al., 2024). Therefore, increasing the length of the HRT can potentially resolve this issue. To test this, editing plasmids targeting *pilB* with either 2kb or 3kb 5’ and 3’ HAs were generated. Both plasmids produced exconjugants, with the 3kb HA plasmid producing slightly more than the 2kb HA plasmid, and both phenotypic and genotypic analyses indicating successful deletion of *pilB* (Fig. 2A-C). We subsequently targeted three additional genes known to be involved in motility, *pilT1*, *pilT2*, and *hmpD*, (Khayatan et al., 2015; Risser et al., 2014; Risser and Meeks, 2013) starting with plasmids containing 2kb HAs. The *pilT2* gene was successfully deleted, but the plasmid targeting *pilT1* failed to produce exconjugants. However, a plasmid targeting *pilT1* and containing 3kb HAs produced mutant exconjugants with the *pilT1* gene deleted (Fig. 2A-C).

In contrast, the plasmid targeting *hmpD* produced a nearly confluent lawn of exconjugants that retained motility, indicating that while the plasmid was successfully introduced, the *hmpD* gene was not deleted. The most likely explanation for this observation is that the gRNA selected for this plasmid is not efficiently directing cleavage of *hmpD*. Previous work has indicated that the chance of selecting a low efficiency gRNA can be overcome by building two separate plasmids with different gRNAs targeting the same GOI (Niu et al., 2019). In our case, because the gRNA array is generated via gene synthesis, this allows for the simultaneous incorporation of two different gRNAs targeting the GOI into the same CRISPR array. A plasmid was subsequently built targeting *hmpD* with 2kb HAs and 2 gRNAs and successfully deleted *hmpD* (Fig. 2A-C).

To further explore the efficacy of this 2-gRNA strategy and the effect of HA length, 8 additional genes known to be required for motility (Hassan et al., 2024; Khayatan et al., 2015; McDonald et al., 2022; Risser et al., 2014) were targeted with plasmids containing 2 gRNAs and HAs ranging from 2-3 kb. For 7 out of the 8 genes, deletion was successful with either 2 or 3 kb HAs (Fig. 2A-C), but for *ebsA*, 4 kb HAs were required for successful gene deletion. Collectively, these results demonstrate that Cpf1-based gene deletion could be implemented in all 14 of the target genes tested, and that incorporation of 2 different gRNAs into the array can decrease the likelihood of inefficient targeting by the gRNA-Cpf1 complex, while larger HAs enhance the probability of recovering mutant exconjugants, likely by enhancing HDR efficiency.

### HIP1 sites do not substantially influence editing efficiency

The genomes of cyanobacteria contain a recurrent motif, Highly Iterative Palindrome 1 (HIP1), with the consensus sequence GCGATCGC (Xu et al., 2018). While the precise function of HIP1 is not entirely defined, recent work has indicated that the presence of HIP1 sites on foreign DNA can enhance the rate of integration via homologous recombination (Kamoku and Nielsen, 2025). Moreover, integration is further enhanced if the HIP1 sites on the foreign DNA are methylated by the methyltransferase from *Synechocystis* sp. PCC 6803 via encoding this enzyme in the genome of an *E. coli* strain used to prepare the foreign DNA (Kamoku and Nielsen, 2025). Therefore, one possible explanation for the variation in the HA length required for successful gene deletion could be related to the presence and number of HIP1 sites on the HRTs. To test this hypothesis, we quantified the number of HIP1 sites present within the HRTs of each plasmid and compared this to the number of exconjugants produced (Fig. 3A). This analysis indicated that there was little correlation between the presence of HIP1 sites and successful gene deletion.

**Figure 3.**
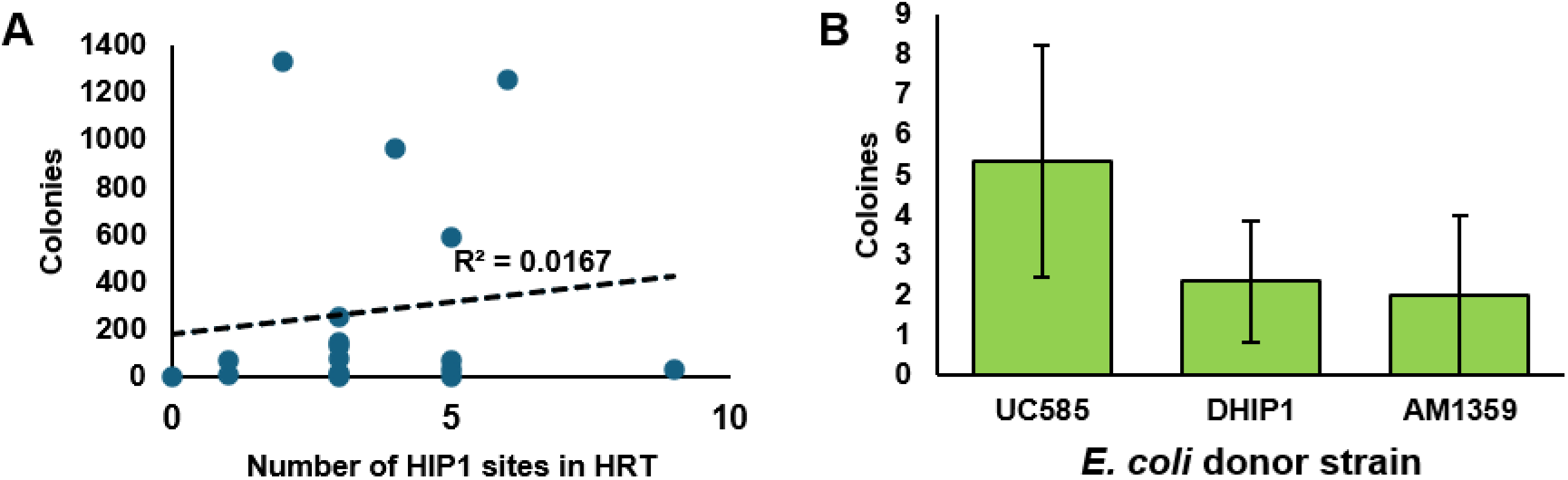
Effect of HIP1 sites on generation of mutant exconjugants. (**A**) Correlation between presence of HIP1 sites in the HRT for an editing plasmid and the average number of exconjugants colonies. (**B**) Average number of exconjugant colonies for the plasmid targeting *pilA* from three independent conjugations using the indicated *E. coli* donor strains. Error bars = +/-1 S.D.

However, it is still possible that methylation of HIP1 sites may improve HDR and thus the efficiency of generating exconjugants. To test this, the helper plasmids pRK24 (Ma et al., 2014b) and pRL528 (Elhai and Wolk, 1988) from UC585 were transformed into *E. coli* strain DHIP1 (Kamoku and Nielsen, 2025), which encodes the HIP1 methyltransferase from *Synechocystis*, and this strain was subsequently used as a donor for conjugal transfer into *N. punctiforme*. The plasmid targeting *pilA* with 2kb HAs was used in this test as it typically yielded only a handful of colonies, indicating that HDR may be limiting for this plasmid, and because it contains a HIP1 site in its 5’ HA. The pCpf1b plasmid backbone also contains a single HIP1 site. However, on average, the UC585 donor strain produced more colonies than the DHIP1 donor strain, indicating that methylation of HIP1 sites in this case did not positively influence conjugation efficiency (Fig. 3B). *E. coli* AM1359 (Ma et al., 2014a), another commonly used donor strain for conjugation with *Nostoc* sp. PCC 7120, which contains the helper plasmids pRL443 and pRL623 (Elhai et al., 1997), was also tested, but did not improve conjugation efficiency compared to UC585 either (Fig. 3B). These results indicate that UC585 appears to be the most efficient donor strain for conjugal transfer into *N. punctiforme*, and, at least in this test case, that methylation of HIP1 sites does not significantly affect conjugation efficiency.

### CRISPR-based allelic replacement

In addition to targeted deletion of genes, CRISPR-based genome editing can be used for allelic exchange, introducing altered alleles with point mutations or various tags. Typically, in addition to generating the desired alterations to the gene of interest, site directed mutagenesis must be performed to remove the PAM site from the target gene so that the gRNA-Cpf1 complex does not cleave the altered allele on the HRT (Ungerer and Pakrasi, 2016). When constructing the plasmids for targeted deletions as described above, the 5’ and 3’ HAs were joined by a common linker. This facilitates more consistent plasmid assembly and results in the introduction of an in-frame exogenous target site in place of most of the gene in the genome (Fig. 4A). Subsequently, altered alleles can be re-integrated at their native loci by introducing a pCpf1b-based plasmid expressing gRNAs that target the exogenous site and contain HRTs with the altered alleles (Fig. 4A). This strategy eliminates the need to incorporate any additional mutations in the GOI to remove PAM sites. As proof of principle, this strategy was tested by constructing a plasmid to re-introduce the *pilA* gene into the Cpf1-generated *pilA* mutant described above. Two different gRNAs targeting the exogenous site were tested (Fig. S4), by constructing gRNA arrays expressing two copies of one or the other of these gRNAs. In both cases, *pilA* was successfully reintegrated at its native locus, based on both genotype analysis and the restoration of motility (Fig. 4 B+C), indicating this can serve as an efficient strategy for generating strains with altered alleles at their native chromosomal locus.

**Figure 4.**
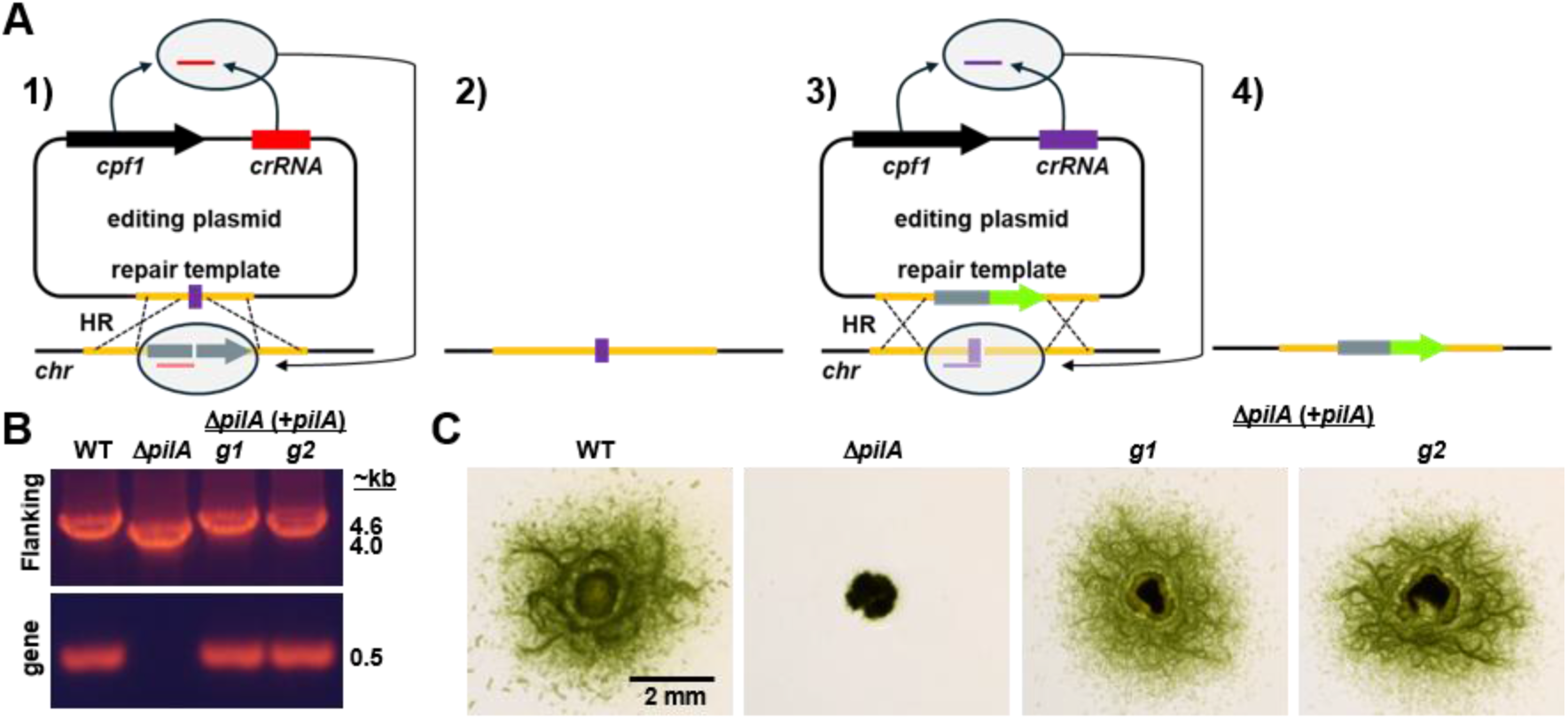
Cpf1-based allelic replacement. (**A**) Schematic diagram of the allelic replacement strategy. 1-2: Construction of deletion mutants results in the integration of an exogenous target site in place of the deleted gene (purple). 3-4: A second plasmid is constructed with a gRNA array targeting the exogenous site and an HRT containing the altered allele and introduced into the previously generated deletion mutant. In this example the altered allele is tagged with *gfp* (green). (**B**) PCR-based genotype analysis of reintroduction of *pilA* at its native chromosomal locus using plasmids containing two copies of one of two different gRNAs (g1 or g2) targeting the exogenous site, as described in Fig. 2B. (**C**) Phenotypic analysis of motility by colony spreading assays.

### Implementation of Cpf-based tools on previously uncharacterized genes

The mutants generated so far in this study have been previously characterized via gene deletion using homologous recombination. To further test the efficacy of these Cpf1-based tools, we subsequently targeted several genes for deletion that have not yet been characterized. The chosen genes encode proteins containing proline-glutamate-proline C-terminal domains (PEP-CTERM) that may play a role in motility. A recent study demonstrated that Cyanoexosortase B (CrtB) is essential for motility in *N. punctiforme* and is thought to recognize and cleave PEP-CTERM domains (Parrett et al., 2025). Seven of the 35 genes encoding PEP-CTERM proteins in *N. punctiforme* are upregulated in developing hormogonia (Gonzalez et al., 2019), and it is likely that the failure to process these proteins in the *crtB* mutant accounts for the defect in motility (Parrett et al., 2025). Because we suspect there may be functional redundancy among this gene set, complex genetic backgrounds containing multiple deletions may need to be generated to elucidate their phenotype, making this an ideal test case for Cpf1-based genetic tools.

To generate deletion mutants, 2kb HAs were initially tested, along with the 2-gRNA strategy, for five of these PEP-CTERM genes. Deletion mutants were successfully generated for Npun_F0296, Npun_F4801, and Npun_R6196 (Fig. 5A+B). However, For Npun_R3960 and Npun_R4127, plasmids with 2kb HAs failed to generate any exconjugants (Fig. 5A). However, when plasmids with 4kb HAs were tested for these two genes, they successfully generated mutants (Fig. 5A+B), which further demonstrates that larger HRTs improve editing efficiency, possibly due to enhancing the rate of HDR. To test the CRISPR-based allelic replacement strategy described above, we attempted to reinsert FLAG-tagged (Hopp et al., 1988) alleles for three of these genes at their native loci. The plasmids utilized HAs of corresponding lengths to those used to generate the deletion mutants. In all three cases, the FLAG-tag allele was successfully inserted into the native locus of the deletion mutants (Fig. 6A+B), which can be used for subsequent immunological studies to investigate localization and processing by CrtB in future studies. Together, these results indicate that the Cpf1-based genetic tools developed in this study can consistently generate deletion mutants and perform allelic replacements.

**Figure 5.**
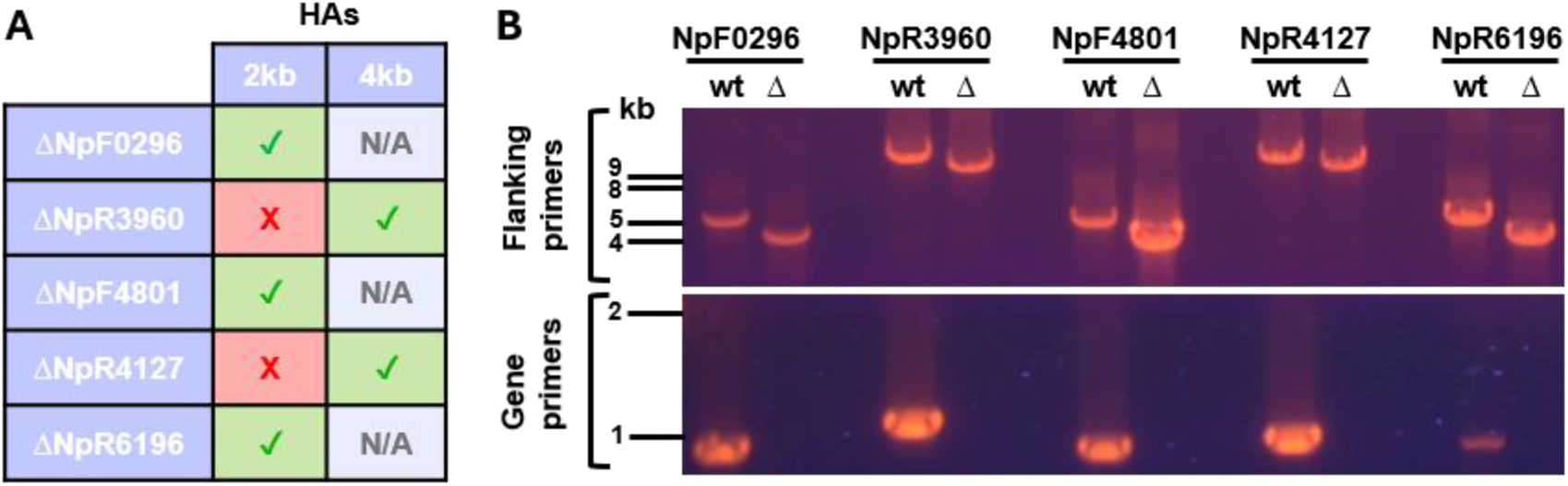
Cpf1-based gene deletion of previously uncharacterized genes. (**A**) Table depicting results for deletion of the indicated genes using editing plasmids with either 2 kb or 4 kb HAs. (**B**) PCR-based genotype analysis of mutants as described in figure 2B.

**Figure 6.**
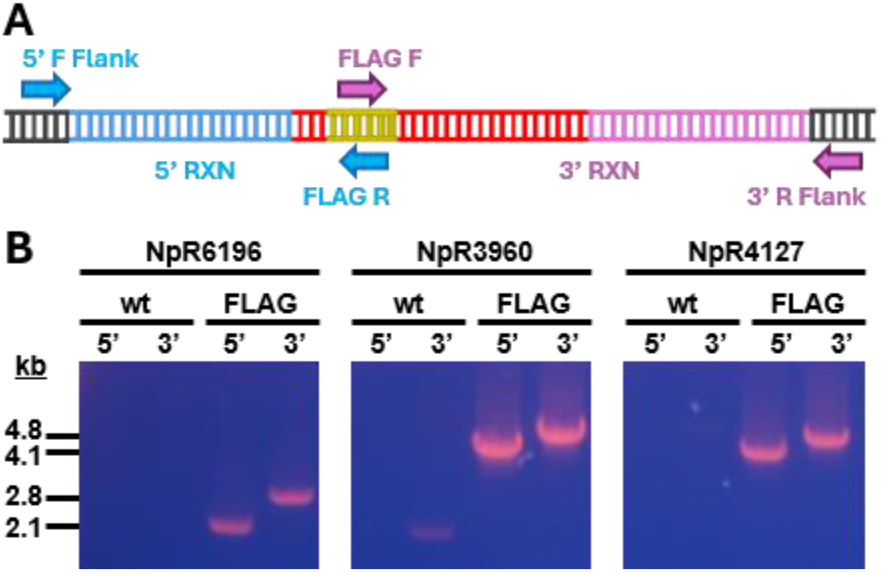
Cpf1-based introduction of FLAG-tagged alleles. (**A**) Schematic diagram of the annealing sites for primers used in PCR reactions to verify the genotype of strains with FLAG-tagged alleles. (**B**) PCR-based genotype analysis of strains with FLAG-tagged alleles. Flanking primers targeting the regions for the GOI (as indicated) and FLAG-specific primers were used in the wild type (wt) or the strain harboring the FLAG-tagged allele (FLAG).

## Discussion

The results from this study demonstrate that Cpf1-based gene deletion and allelic replacement can be consistently implemented in *N. punctiforme*. This approach was also substantially faster than mutant generation via homologous recombination, generating mutant strains in ∼2 months or less, including curing of the editing plasmid, while homologous recombination typically takes ∼4 months under ideal circumstances. It was also considerably less labor intensive than homologous recombination because nearly all the exconjugant colonies were mutants and thus did not require extensive PCR-based screening to identify positive clones.

One critical finding for successful mutant generation in *N. punctiforme* using Cpf1 is the variation in HRT length requirement. While 4 kb HAs were the largest required for deletion of any of the genes tested in this study, it is possible that there are other genes that may require even larger HRTs. Notably, with the cloning strategy implemented in this study we did not encounter any issues constructing plasmids with larger HRTs, and the wide availability of inexpensive nanopore-based sequencing (MacKenzie and Argyropoulos, 2023) allowed us to easily ensure sequence fidelity even of large constructs. Thus, we do not envision construction of editing plasmids with large HRTs to be a significant barrier to implementing this approach. Based on the data presented, we would suggest that ∼2-4 kb HAs are an ideal starting point for targeting novel genes. However, it is worth noting that in some cases, larger HAs may be detrimental. For instance, they might lead to the introduction of genes that are deleterious at higher gene dosage. Therefore, the size of the HRT used may need to be adjusted to some degree on a case-by-case basis.

The underlying mechanism for why some genes require larger HRTs for deletion is currently unclear. As demonstrated in this work, it does not appear to be related to the presence of HIP1 sites in the foreign DNA, nor the methylation status of these sites. There was also no obvious correlation with gene size. Moreover, when using homologous recombination, we have always found ∼1kb HAs to be sufficient for successful integration of suicide vectors into the genome, indicating that the HRT length requirement is unique to Cpf1-based editing and possibly HDR. One possibility is that the HRT length requirement may be related to the efficiency of targeted cleavage by the gRNA. It may be that highly efficient gRNAs will rapidly cleave all copies of the chromosome upon expression in a recipient cell, leading to cell death before HDR can repair the lesions. In this case, longer HRTs may enhance the rate of HDR to compensate for the rapid rate of cleavage. Further work would be needed, testing many different gRNAs targeting the same gene, and using HRTs of various length for HDR, to test this theory. To our knowledge, this requirement for larger HRTs has not been observed in other cyanobacteria where Cpf1-based editing has been implemented, but it is possible that this may pose a barrier in other non-traditional model cyanobacteria.

We envision these tools will be particularly powerful for rapid generation of multiple mutations in a single strain to test for functional redundancy, as discussed for the PEP-CTERM genes targeted in this study, or for studies such as epistasis analysis. Moreover, the protocols developed and described here should allow for easy implementation of Cpf1-based genetic manipulation by other laboratories, enhancing the pace of genetic studies to investigate various aspects of *N. punctiforme* biology.

## Materials and Methods

### Strains and culture conditions

*N. punctiforme* ATCC 29133 and its derivatives were cultured in Allen and Arnon medium diluted four-fold (AA/4) (Allen and Arnon, 1955) (https://www.protocols.io/workspaces/risser-lab-n.punctiforme) under constant white light illumination (∼50 μmol photons m^−2^ s^−1^) at room temperature (∼25°C) in ambient atmosphere without any additional carbon supplementation (*e.g.* sodium bicarbonate). Unless otherwise stated, cultures were also grown without supplementation of fixed nitrogen. For liquid cultures, 50 mL of AA/4 in 125 mL Erlenmeyer flasks were inoculated and incubated under illumination with shaking at 120 rpm. Solid medium contained 1% w/v noble agar. For selective growth, the medium was supplemented with 30-50 μg ml^-1^ neomycin as indicated. *Escherichia coli* cultures were grown in lysogeny broth (LB) for liquid cultures or LB supplemented with 1.5% (w/v) bacto agar for plates. Selective growth medium was supplemented with 50 μg ml^-1^ kanamycin, 50 μg ml^-1^ ampicillin, and 15 μg ml^-1^ chloramphenicol.

### Plasmid construction

For a detailed description of plasmids and primers used in this study, refer to table S1. Detailed protocols for Cpf1-based plasmid construction and gene deletion can also be found at (https://www.protocols.io/workspaces/risser-lab-n.-punctiforme). To construct plasmids for Cpf1-based in-frame deletions of target genes, 5’ and 3’ HAs of varying lengths between ∼1-4 kb (depending on the target gene) of flanking DNA on either side of the gene and several codons at the beginning and end of the gene were amplified via PCR. In a total reaction volume of 50 μL, 1 μL of 100 ng/μL *N. punctiforme* chromosomal DNA was used as template, along with 25 μL of Platinum SuperFi II PCR Master Mix (ThermoFisher Scientific) and 0.5 μL of each primer (50 μM stock concentration), following the manufacturer’s recommendations for the thermocycler conditions. Gene synthesis (GenScript) was used to generate gRNA arrays replacing either the first, or both the first and second spacers with 22 bp gRNAs targeting the gene of interest adjacent to a TTTG PAM site, cloned into pUC57. These gRNA arrays were then amplified via PCR from the plasmids. In a total of 50 μL, 1 μL of ∼4 ng/μL plasmid containing the gene-specific gRNA array was used as template with 25 μL Platinum SuperFi II PCR Master Mix and 0.5 μL of the primers (50 μM stock) gRNA-array-F-AvrII and pCpf1b-rep/trm-R following the manufacturer’s recommendations for thermocycler conditions. Subsequently, 1 μL of each unpurified PCR product was combined with ∼80 ng of column purified pCpf1b (Niu et al., 2019), digested with AvrII and BamHI, and 10 μL of HiFi assembly master mix (New England Biolabs, Inc.) in a total volume of 20 μL and incubated for 1 h at 50°C. Two μL of the assembly reaction were transformed into NEB 5-α competent cells (New England Biolabs, Inc.) following the manufacturer’s instructions, and plated on LB containing 50 μg/mL kanamycin, 20 μg/mL X-gal, 0.1 mM IPTG. White colonies were picked to inoculate overnight liquid cultures, and the plasmids were subsequently purified using the Monarch Plasmid Miniprep Kit (New England Biolabs, Inc.) and sequenced (Plasmidsaurus) to ensure fidelity.

Construction of plasmids for reintegration of altered alleles at their native chromosomal was performed as described for deletion plasmids, except that the template used for amplification of the gRNA array contained two copies of a gRNA targeting the exogenous target site integrated into the genome in place of deleted genes. For the plasmids used to reintegrate *pilA* at its native chromosomal locus, the homologous repair template generated by PCR contained the coding sequence and ∼2kb of upstream and downstream DNA. For the plasmids used to reintegrate FLAG-tagged allele, the homologous repair template was generated by PCR in two separate reactions. The first reaction amplified ∼2-4 kb (depending on the target gene) of DNA upstream of the gene, and the codons corresponding to the signal peptide, followed by codons for the FLAG-tag, which were introduced on the primer used for amplification. The second reaction amplified the codons for the FLAG-tag, which were introduced on the primer used for amplification, followed by the remainder of the coding region and ∼2-4kb (depending on the target gene) downstream of the gene. The FLAG-tag codons served as the complementary ends of the 5’ and 3’ HAs for subsequent HiFi assembly.

### Strain construction

To prepare *N. punctiforme* for conjugation, liquid cultures were grown for ∼4-weeks in liquid AA/4 medium supplemented with 3 mM sucralose to prevent hormogonium development(Splitt and Risser, 2016), and pipetted weekly to produce well dispersed cultures amenable to conjugation. The chlorophyll *a* (Chl *a*) concentration was determined and cultures used for conjugation typically had a Chl *a* concentration of between 12.5-25 μg/ml. The cultures were then concentrated by centrifugation at 2,000 rcf for 3 min and removal of the appropriate volume of supernatant to yield a final concentration of 62.5 μg Chl *a*/mL. The cell pellets were then vigorously resuspended by pipetting in this final volume.

To prepare *E. coli* for conjugation, 2 μL of purified plasmid DNA was transformed into *E. coli* UC585 (Liang et al., 1993) and plated onto LB plates containing kanamycin (50 μg/mL), ampicillin (50 μg/mL), and chloramphenicol (15 μg/mL). A single colony was picked from the transformation plate to inoculate 20 mL LB with kanamycin (50 μg/mL), ampicillin (50 μg/mL), and chloramphenicol (15 μg/mL) in a 50 mL conical tube. Caps were cross threaded to allow gas exchange and incubated overnight at 37°C with shaking at ∼250 RPM until an OD_600_ ∼ 1.5-2 was reached. Cultures were centrifuged at 3,234 rcf for 10 min, the supernatant decanted, and the pellet resuspended in 4 mL LB (without antibiotics). The cells were centrifuged again at 3,234 rcf for 10 min, the supernatant decanted, and the cells resuspended in 400 μL LB (without antibiotics). The 400 μL of *E. coli* cells were then combined with 400 μL of the concentrated *N. punctiforme* culture described above and mixed by pipetting. 400 μL of the *N. punctiforme*/*E. coli* cell mixture was plated onto each of two AA/4 plates (containing 30 mL of medium) supplemented with 2.5 mM NH_4_Cl and 5 mM MOPS (pH 7.8). Plates were incubated 24 h at 37°C in the dark and then incubated in the light at room temperature (∼25°C) (lid side down) for 24 h. 10 mL of AA/4 liquid medium with neomycin (30 μg/mL) was then pipetted onto the surface of each plate. Plates were incubated (lid side up) for 2-3 days before removing condensation that accumulated on the lid. Condensation was then removed daily until each plate was dry, and then each plate was sealed with electrical tape and incubated (lid side down) for 2-3 weeks until colonies appeared.

Neomycin resistant colonies were picked and streaked onto fresh AA/4 plates supplemented with 2.5 mM NH_4_Cl, 5 mM MOPS, and neomycin (50 μg/mL) and screened via colony PCR with primers that anneal to the target gene to identify deletion mutants. To cure the plasmid, positive clones were transferred to AA/4 liquid cultures without antibiotic selection, and supplemented with 2.5 mM NH_4_Cl, 5 mM MOPS (pH 7.8), and 3 mM sucralose to suppress hormogonia development and promote dispersed cultures. The cultures were incubated for 2 weeks under illumination with shaking at 120 rpm. Subsequently, 1 mL of culture was transferred to a 1.5 mL tube and vigorously pipetted to disperse the filaments, after which a 1:100 dilution was prepared and 400 μl of the dilution was plated on AA/4 supplemented with 2.5 mM NH_4_Cl, 5 mM MOPS pH 7.8 and 5% w/v sucrose to select for clones that had cured the pCpf1b plasmid. These were subsequently checked for neomycin sensitivity to confirm the loss of the plasmid.

### Genomic DNA isolation

To prepare purified DNA for final PCR verification of mutant strains, chromosomal DNA isolation was performed using the QIAmp Mini Kit (Qiagen). *N. punctiforme* pellets were suspended in 180 μL of lysozyme solution (20mg/mL lysozyme; 20mM tris-HCl, pH 8.0; 2mM EDTA; 1.2% Triton) and mixed by gently pipetting up and down. Mixed pellets were then incubated for 20 minutes at 37°C. After incubating for 20 minutes, 200 μL lysis buffer and 20uL proteinase K solution were added. The samples were then vortexed for 15 seconds and subsequently incubated at 56°C for 20 minutes and then for 15 minutes at 95°C. After incubation, the tubes were centrifuged at 16,000 x g for one minute. The supernatant was transferred to a new tube, 200 μL of 100% ethanol was added and the sample vortexed for 15 seconds. This mixture was then added to a spin column from the QIAmp Mini Kit and subsequent centrifugation and wash steps performed following the manufacturer’s instructions. The genomic DNA was eluted in a final volume of 50 μl elution buffer.

### PCR verification of mutant strains

For a detailed description of the primers used to verify mutant strains, refer to table S1. To verify the genotype of deletion mutant strains, chromosomal DNA (∼100 μg) was used as template in PCR reactions with primers that flanked the region used for the HRT, or annealed to the beginning and end of the deleted gene. To verify the integration of FLAG-tagged alleles, primers that flanked the region used for the HRT and primers that annealed to the codons for the FLAG epitope were used. Amplicons less than ∼2kb were generated using MangoMix (Bioline), while those larger than ∼2kb were generated using RangerMix (Bioline), following the manufacturer’s instructions.

### Phenotypic analysis of motility

Colony motility assays were used to verify motility deficiency in mutant strains. Strains were first streaked on AA/4 5% sucrose plates to suppress hormogonium development. Isolated colonies were subsequently transferred to AA/4 0.5% noble agar and incubated for 5 days before imaging with a Leica SD9 dissecting microscope equipped with a Leica Flexcam C3 camera controlled by Leica LAS X software.

## Acknowledgements

This work was supported by NSF-IOS 2420339 (D.D.R). *E. coli* DHIP1 was a gift from David Nielsen (Addgene plasmid # 226851) (Kamoku and Nielsen, 2025). *E. coli* AM1359 (Ma et al., 2014a) was a gift from James Golden.

