## Supplemental material for "Cpf1(Cas12a)-based genome editing in the filamentous cyanobacterium *Nostoc punctiforme*"

### *pilB (Npun_R0118)-1x-gRNA*

ctagcgctgatttaggcaaaaacgggtctaagaactttaaataatttctactgttgtagat
GATAACTCCTCCACCCAACTGGgtctaagaactttaaataatttctactgt
tgtagatgctactattcctgtgccttcagataattcagtctaagaactttaaataatttctact
gttgtagatgtctagagccttttgtattagtagccggtctaagaactttaaataatttctact
gttgtagattagcgatttatgaaggtcatttttttgtctagctttaatgcggtagttggtaccgt
cgatggcagaaattcgatatctagatctcat

### *hmpB (Npun_F5961)-2x-gRNA*

ctagcgctgatttaggcaaaaacgggtctaagaactttaaataatttctactgttgtagat
TCGGCGGTTAAAATCCGATCCAgtctaagaactttaaataatttctactgt
tgtagatATCGTTACTGGGGTATGAAACAgtctaagaactttaaataattt
ctactgttgtagatgtctagagccttttgtattagtagccggtctaagaactttaaataattt
ctactgttgtagattagcgatttatgaaggtcatttttttgtctagctttaatgcggtagttggt
accgtcgatggcagaaattcgatatctagatctcat

**Figure S1.** Sequence of synthesized gRNA arrays. Example sequences of gRNA arrays containing either 1 gRNA or 2 gRNAs are depicted. The gRNAs are highlighted in red.

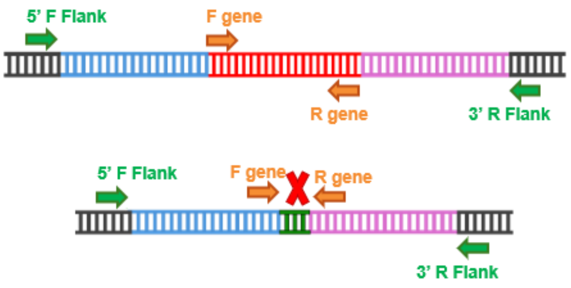


**Figure S2.** Schematic diagram of primer binding sites for primers used in PCR reactions to verify genotype of the mutant strains. 5' F Flank and 3' R Flank anneal just outside of the region used for the HRT and should produce a smaller PCR product if the gene is deleted, while F gene and R gene anneal to the beginning and end of the GOI and should fail to produce a PCR product if the gene is deleted.

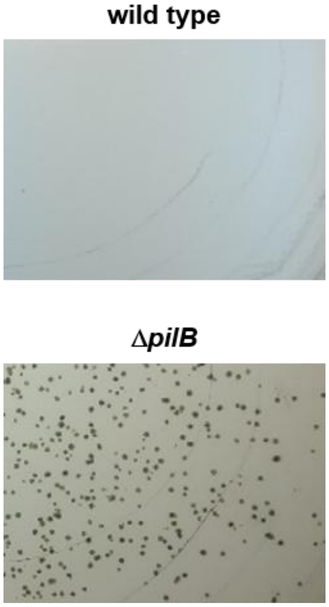


**Figure S3.** Images for sections of plates following introduction of the editing plasmid targeting *pilB*, with a single gRNA and 1 kb HAs, into the wild type or Δ*pilB* strain.

**Exo-F-2x-gRNA (g1)**

GTTTGGAGAAGTCATTTAATAAGGCCACTGTTAAAA

**Exo-R-2x-gRNA (g2)**

GTTTGGAGAAGTCATTTAATAAGGCCACTGTTAAAA

**Figure S4.** Sequence for the exogenous target site integrated into the genome in place of deleted gene with the PAM sites (underlined) and gRNA sequence (Yellow Highlight) for the two gRNA sequences targeting this site.
